# The effect of melittin intervention on murine cervical cancer cells: An in-depth proteomics investigation

**DOI:** 10.64898/2026.08.02.742264

**Authors:** Ronghua Zhang, Mengyi Wang, Kaiyao Zhang, Haiwen Zhuo, Shaode Li, Jianrong Jiang, Jianfeng Qiu, Dafu Chen, Tizhen Yan, Rui Guo

## Abstract

Melittin, the principal bioactive peptide of bee venom, exhibits promising antitumor activity, whereas its molecular mechanisms in cervical cancer remain incompletely understood. In this study, the biological effects and molecular responses of melittin in U14 cervical cancer cells were investigated using Astral data-independent acquisition (Astral-DIA)-based quantitative proteomics combined with molecular validation. The effects of melittin on cell migration, invasion, and cell death were evaluated by Transwell assays and PI/Hoechst staining. Differentially expressed proteins (DEPs) were screened and subjected to Gene Ontology (GO), Kyoto Encyclopedia of Genes and Genomes (KEGG), and protein–protein interaction (PPI) analyses. Representative oxidative stress-related genes and proteins were further validated by RT-qPCR and Western blotting. Melittin significantly inhibited the migration and invasion of U14 cervical cancer cells and increased cell death. Quantitative proteomics identified 9,782 protein groups and 187 DEPs, including 71 up- and 116 down-regulated proteins. KEGG pathway enrichment analysis revealed oxidative phosphorylation (OXPHOS) as the most significantly enriched pathway, together with glutathione metabolism, ferroptosis-related pathways, reactive oxygen species signaling, and mitophagy. GO term enrichment analysis indicated that DEGs were mainly engaged in mitochondrial function, electron transport, oxidoreductase activity, and energy metabolism. RT-qPCR assay demonstrated altered expression of *Duox*1, *Gpx*4, *Gsx*2, *Nfe*2l2, and *Gstp*2. Additionally, PPI analysis identified *Gstp*2 and ODC1 as representative hub proteins involved in redox regulation and metabolic adaptation. Furthermore, Western blotting confirmed increased GSTP2 expression following melittin treatment. Overall, these findings provide a comprehensive proteomic landscape of melittin-treated U14 cervical cancer cells and suggest that mitochondrial OXPHOS remodeling and redox-associated pathways may contribute to the antitumor activity of melittin.

## 1. Introduction

Melittin is the principal bioactive peptide of bee venom, accounting for approximately 45–50% of its dry weight. It is a linear cationic amphipathic peptide consisting of 26 amino acid residues, with a molecular weight of 2.83 kDa and a molecular formula of C₁₃₁H₂₂₉N₃₉O₃₁ [1,2]. Owing to its unique amphipathic α-helical structure, melittin exhibits a wide range of biological activities, such as anti-inflammatory, antimicrobial, immuno-modulatory, neuroprotective, and antitumor effects [3,4]. Traditionally, the biological activity of melittin has been primarily attributed to its interaction with negatively charged phospholipid membranes. Following membrane binding, melittin inserts into the lipid bilayer and forms transmembrane pores, increases membrane permeability, and induces leakage of intracellular components, ultimately causing cell lysis [5–8]. This membrane-disruptive property contributes substantially to its antimicrobial and antitumor activities, however, it limits the clinical application of melittin because of cytotoxicity toward normal cells [9]. Consequently, numerous studies have focused on improving the tumor selectivity and therapeutic safety of melittin through structural modification and targeted delivery strategies [10–13].

Beyond its membrane-disruptive activity, accumulating evidence suggests that melittin also exerts profound intracellular regulatory effects. Previous studies have demonstrated that melittin suppresses tumor cell proliferation by inducing oxidative stress, mitochondrial dysfunction, apoptosis, autophagy, ferroptosis, and cell-cycle arrest, while simultaneously regulating multiple signaling pathways, including PI3K/Akt, MAPK, NF-κB, and JAK/STAT [14–20]. Nevertheless, these studies have mainly focused on individual signaling pathways or specific biological processes. The global alterations in protein expression and molecular regulatory networks triggered by melittin remain largely unexplored. Therefore, a comprehensive proteomic analysis is required to systematically characterize the molecular responses induced by melittin and to identify novel therapeutic targets.

Cervical cancer is the fourth most common malignancy and the fourth leading cause of cancer-related death among women worldwide [21,22]. Although surgery, radiotherapy, chemotherapy, and targeted therapies have significantly improved patient outcomes, treatment failure caused by drug resistance, recurrence, and systemic toxicity continues to pose substantial clinical challenges [23]. Increasing evidence indicates that cervical carcinogenesis is accompanied by extensive metabolic reprogramming, dysregulated redox homeostasis, and aberrant protein expression [24,25]. Consequently, identification of key proteins and signaling pathways involved in cervical cancer progression is essential for elucidating disease mechanisms and developing more effective therapeutic strategies. Proteomics, particularly DIA-based quantitative proteomics, has emerged as a powerful weapon for comprehensive and highly reproducible protein profiling. Compared with transcriptomic analysis, proteomics directly reflects changes in protein abundance and biological function, enabling the systematic identification of differentially expressed proteins (DEPs), dysregulated signaling pathways, and potential therapeutic targets [26,27]. Indeed, proteomic approaches have successfully been applied to investigate the molecular mechanisms of natural products and bioactive peptides, including melittin, in diverse biological systems [28,29].

In a recent study, we obtained high-quality transcriptome data of melittin- and untreated cervical cancer U14 cervical cancer cells [30], based on which we analyzed the effect of melittin intervention on U14 cervical cancer cells at transcriptome level [31]. Despite the growing interest in melittin as a promising anticancer peptide, its effect on global proteome and underlying molecular mechanisms in cervical cancer have not yet been systematically characterized. Here, U14 cervical cancer cells were treated with melittin at different concentration, followed by Cell apoptosis was assessed by PI/Hoechst double staining, and positively stained cells were observed and counted under a fluorescence microscope. Additionally, the melittin- and un-treated U14 cell samples were subjected to deep proteomic profiling utilizing Astral-DIA-based quantitative proteomics, and DEPs were identified and subsequently analyzed to elucidate the biological processes and signaling pathways regulated by melittin. Moreover, reverse transcription quantitative PCR (RT-qPCR) and Western blot were combined to perform verification of randomly selected DEPs. This study not only reveals the effect of melittin on the invasion, migration, and cell viability of cervical cancer cells, but also offers a comprehensive proteomic landscape of melittin-intervened cervical cancer cells. Our findings could identify potential molecular targets and regulatory pathways associated with its antitumor activity, thus paving a path for development and clinical application of melittin as an anticancer agent in the future.

## 2. Materials and Methods

### 2.1 Cell culture

The murine cervical carcinoma cell line U14 was purchased from Xiamen Yimao Biotechnology Co., Ltd. (Xiamen, China). Cells were cultured in high-glucose Dulbecco’s Modified Eagle Medium (DMEM; Gibco, USA) supplemented with 10% fetal bovine serum (FBS; Gibco, USA) and 1% penicillin–streptomycin solution (Yeasen, China). Cells were maintained at 37°C in a humidified incubator with 5% CO₂ and passaged every 2 days. Cells in the logarithmic growth phase were used for subsequent experiments.

### 2.2 Melittin preparation

Melittin (purity >99%) was purchased from Selleck Chemicals (Houston, TX, USA). The lyophilized peptide was dissolved in sterile water to prepare a 4 mg/mL stock solution, aliquoted, and stored at −80°C protected from light. Before use, the stock solution was diluted with complete DMEM to the desired working concentrations.

### 2.3 Transwell Invasion Assay

U14 cervical cancer cells were harvested after 24 h of culture and resuspended in serum-free medium. A total of 200 µL of the cell suspension was added to the upper chamber of a Transwell insert pre-coated with 50 µL of Matrigel, which was then incubated at 37°C for 1 h to allow gel polymerization. After the Matrigel had solidified, the excess Matrigel was carefully removed. The upper chamber was equilibrated with complete medium to prevent leakage before the prepared cell suspension was gently added. The lower chamber was filled with culture medium supplemented with 20% FBS as a chemoattractant. After 24 h of incubation, the inserts were washed with 0.01 M phosphate-buffered saline (PBS), fixed with 4% paraformaldehyde for 20 min, and then stained with 0.1% crystal violet for 15 min following a second PBS wash. Non-invading cells on the upper surface of the membrane were gently removed using a cotton swab. Invaded cells on the lower surface of the membrane were visualized under a microscope, and the mean number of invaded cells was quantified.

### 2.4 Cell Sample Preparation

Cells were removed from the incubator, followed by aspiration of the culture medium. The cells were washed three times with pre-chilled 0.01 M PBS to remove residual medium. Cells in the treatment group were then incubated with 4 µg/mL melittin solution, ensuring complete coverage of the cell monolayer, and then maintained at 37°C in a humidified incubator containing 5% CO₂ for 20 min. Subsequently, the culture dishes were placed on ice and the cells were washed with pre-chilled 0.01 M PBS. The cells were gently scraped from the culture dish with a cell scraper and collected into sterile centrifuge tubes. Following centrifugation, the cell pellets were transferred to pre-chilled cryogenic tubes for subsequent experiments.

### 2.5 Assessment of cell viability and membrane integrity

U14 cervical cancer cells were seeded at a density of 1 × 10⁴ cells per well in 96-well flat-bottom plates and allowed to adhere overnight. U14 cervical cancer cells were randomly assigned to a melittin-treated group (T) and an untreated control group (C). Following exposure to graded concentrations of melittin (0, 2, 4, 6, and 8 µg/mL) for 20 min, cells were co-stained with Hoechst 33342 (10 µg/mL) and propidium iodide (PI, 5 µg/mL) in the dark at 37°C for 15 min. After three washes with PBS, fluorescent images were acquired at 200× magnification using an inverted fluorescence microscope (Nikon, Tokyo, Japan). For each well, at least five non-overlapping fields were randomly selected, and both Hoechst-positive (total nuclei) and PI-positive (membrane-compromised nuclei) cells were enumerated using ImageJ software [32]. The percentage of PI-positive cells was normalized to the total Hoechst-positive nuclei and expressed as the PI/Hoechst ratio. Data were pooled from three independent experiments, each performed in triplicate.

### 2.6 RT-qPCR assay

To further investigate the molecular mechanisms, RT-qPCR was performed under the same treatment conditions. Total RNA was extracted from U14 cervical cancer cells using the SteadyPure RNA Extraction Kit (Accurate Biology, Hunan, China) according to the manufacturer’s instructions. RNA concentration and purity were determined using a NanoDrop™ 2000 spectrophotometer (Thermo Fisher Scientific, USA), and samples with an A₂₆₀/A₂₈₀ ratio between 1.8 and 2.0 were processed for reverse transcription. Complementary DNA (cDNA) was synthesized from 1 µg of total RNA using the Hifair® AdvanceFast 1st Strand cDNA Synthesis Kit (Yeasen, Shanghai, China). Subsequently, qPCR was carried out with the Hieff® qPCR SYBR Green Master Mix (Yeasen) on a Bio-Rad CFX96™ real-time detection system. The thermal cycling protocol consisted of initial denaturation at 95°C for 30 s, followed by 40 cycles of 95°C for 5 s and 60°C for 30 s. GAPDH served as the internal reference gene, and relative gene expression was determined using the 2 ^ΔΔCt^ method. All reactions were run in triplicate, and three independent biological replicates were performed.

### 2.7 Protein Extraction, Enzymatic Digestion, and Mass Spectrometric Analysis

An appropriate amount of cell sample was weighed and transferred into a 2 mL centrifuge tube, and steel balls, lysis buffer containing 8 M urea/50 mM Tris-HCl, and Roche cOmplete protease inhibitor cocktail (final concentration 1×) were then added. Next, the mixture was placed on ice for 5 min. Samples were homogenized using a tissue lyser (60 Hz, 2 min), followed by centrifugation at 20,000 g for 15 min at 4°C. The supernatant was collected, and DTT was added to a final concentration of 10 mM. The samples were incubated in a 37°C water bath for 1 h. Finally, IAA was added to a final concentration of 20 mM, and the samples were then incubated in the dark for 30 min.

Protein quantification was performed using the Bradford assay. Standard protein solutions (0.2 µg/µL BSA) of 0, 2, 4, 6, 8, 10, 12, 14, 16, and 18 µL were added sequentially into the wells of a 96-well plate, followed by the addition of 20, 18, 16, 14, 12, 10, 8, 6, 4, and 2 µL of pure water, respectively. After mixing thoroughly, 180 µL of Coomassie Brilliant Blue G-250 working solution was added to each well. The absorbance at 595 nm (OD595) was measured utilizing a microplate reader, and a linear standard curve was generated based on OD595 and protein concentration. The test protein solution was appropriately diluted, and 180 µL of the working solution was added to 20 µL of the protein solution. OD595 of the sample was read, and the protein concentration was calculated according to the standard curve and sample OD595. For each sample, 10 µg of protein solution was mixed with an appropriate volume of loading buffer, heated at 95°C for 5 min, and then centrifuged at 20,000 g for 5 min. The supernatant was loaded into the sample wells of a 4–12% SDS polyacrylamide gel and subjected to electrophoresis at a constant voltage of 80 V for 20 min and then 120 V for 60 min. After electrophoresis, the gel was stained, destained, and photographed.

For each sample, 150 µg of protein was taken and digested with 3 µg of Trypsin at a protein:enzyme ratio of 50:1 for 14–16 h at 37°C. The digested peptides were desalted using Waters solid-phase extraction cartridges, vacuum-dried, redissolved in pure water, and stored at –20°C. Equal amounts of peptides from each sample were mixed, diluted with mobile phase A (5% ACN, pH 9.8), and injected. Separation was performed on a Thermo Scientific UltiMate™ 3000 Binary Rapid Separation System using a 3.5 µm 4.6 × 150 mm Agilent ZORBAX 300Extend-C18 column. Gradient elution was carried out at a flow rate of 0.3 mL/min: 5% to 21.5% mobile phase B (97% ACN, pH 9.8) for 38 min, 21.5% to 40% mobile phase B for 20 min, 40% to 90% mobile phase B for 2 min, 90% mobile phase B for 3 min, and 5% mobile phase B for equilibration for 10 min. Elution peaks were monitored at 214 nm, and fractions were collected every minute. Ten fractions were obtained by combining samples according to the chromatographic elution peaks and then freeze-dried. Mobile phases A (100% water, 0.1% formic acid) and B (80% acetonitrile) were prepared. The dried peptide samples were reconstituted with 0.1% formic acid, centrifuged at 20,000 g for 10 min, and the supernatant was injected. Separation was performed using a Thermo Scientific Vanquish Neo UHPLC system. Proteomic analysis was performed by LC-Bio Technology Co., Ltd. (Hangzhou, China) using the Astral-DIA quantitative proteomics platform.

### 2.8 Protein Identification and Quantification

Protein annotation was performed against the UniProt Mus musculus reference proteome (Taxonomy ID: 10090). DIA-NN software was employed for protein search, identification, and quantification of the DIA mass spectrometry (MS) data. The raw MS files were searched against the corresponding database https://www.uniprot.org/uni-protkb?query=organism_id:10090. Protein identification was conducted based on the search results, and peptide and protein distribution analyses were performed to assess the quality of the MS data and database search. Functional annotations of the identified proteins were conducted using four common databases, including GO, KEGG, Reactome, and subcellular localization. Then, protein quantification, sample correlation analysis, and differential expression analysis were performed. For the DEPs, comprehensive functional investigation were conducted, including GO and KEGG enrichment analysis, protein-protein interaction network analysis, and protein correlation analysis.

### 2.9 Western blot

Total protein was extracted from U14 cervical cancer cells using RIPA lysis buffer (yeasen, China) supplemented with Nox Protease Inhibitor Cocktail for Mammalian Cell (G-Biosciences, Cat. 786-387, used at 1:100). Protein concentrations were determined using the BCA assay (epizyme, China). Equal amounts of protein (20 µg) were separated by SDS-PAGE and transferred onto PVDF membranes (Millipore, USA). After blocking with 5% non-fat milk for 2 h at room temperature, membranes were incubated overnight at 4°C with primary antibodies against GSTP2 (Cat. No. ARP93219_P050, Aviva Systems Biology, CA, USA, 1:1000), ODC1 (bs-1294R, Bioss Inc., Woburn, MA, USA, 1:1000, and β-tubulin (Cat. No.66240-1-Ig, Proteintech, Wuhan, China, 1:10000). Following HRP-conjugated secondary antibody incubation, protein bands were visualized using an ECL detection system (Absin Bioscience Inc. Shanghai, China) and imaged with a ChemiDoc™ imaging system (Bio-Rad, USA). Band intensities were quantified using ImageJ software (v1.53, NIH, USA), and target protein levels were normalized to β-tubulin. All experiments were performed in triplicate.

### 2.10 Statistical analysis

Statistical analysis of the data was mainly done by R software (version 4.0). Raw intensity values of proteins were normalized to standardized data by median normalization and minimum complementary values. Statistical *P* values between two groups were obtained by t-test and statistical analysis between multiple groups was performed by one-way ANOVA. Only proteins with a *P* value < 0.05 and a multiplicity of differences > 1.2 were defined as final significant proteins. To identify biological pathways significantly associated with the gene set, we performed gene set enrichment analysis (GSEA) using the Molecular Signatures Database (MSigDB). Enrichment was assessed across multiple functional annotation categories, including Gene Ontology (GO) terms, Kyoto Encyclopedia of Genes and Genomes (KEGG) pathways, Disease Ontology (DO) terms, and Reactome pathways. A pathway was considered significantly enriched if it met the following criteria |NES| > 1, nominal *P* value < 0.05, and false discovery rate (FDR) q value < 0.25. Subcellular localization analysis was performed by WoLF PSORT. PPI analysis was based on the search in the STRING database, and all protein interactions with confidence scores ≥0.4 were extracted for mapping.

## 3. Results

### 3.1 Melittin inhibits the migration and invasion of U14 cervical cancer cells

Transwell migration and invasion assays showed that the number of cells that migrated through or invaded across the Transwell membrane gradually reduced with increasing concentrations of melittin (Figure 1A). At the highest concentration (8 µg/mL), only a few migrated or invaded cells were observed (Figure 1A). In addition, as compared with the un-treated group, melittin (2, 4, 6, and 8 µg/mL) significantly decreased the migration and invasion of U14 cervical cancer cells in a dose-dependent manner, as shown in Figure 1B. Notably, treatment with 8 µg/mL of melittin almost completely abolished the migratory and invasive capacities of U14 cervical cancer cells. These results indicate that melittin markedly inhibits the migration and invasion of U14 cervical cancer cells.

**Figure 1.**
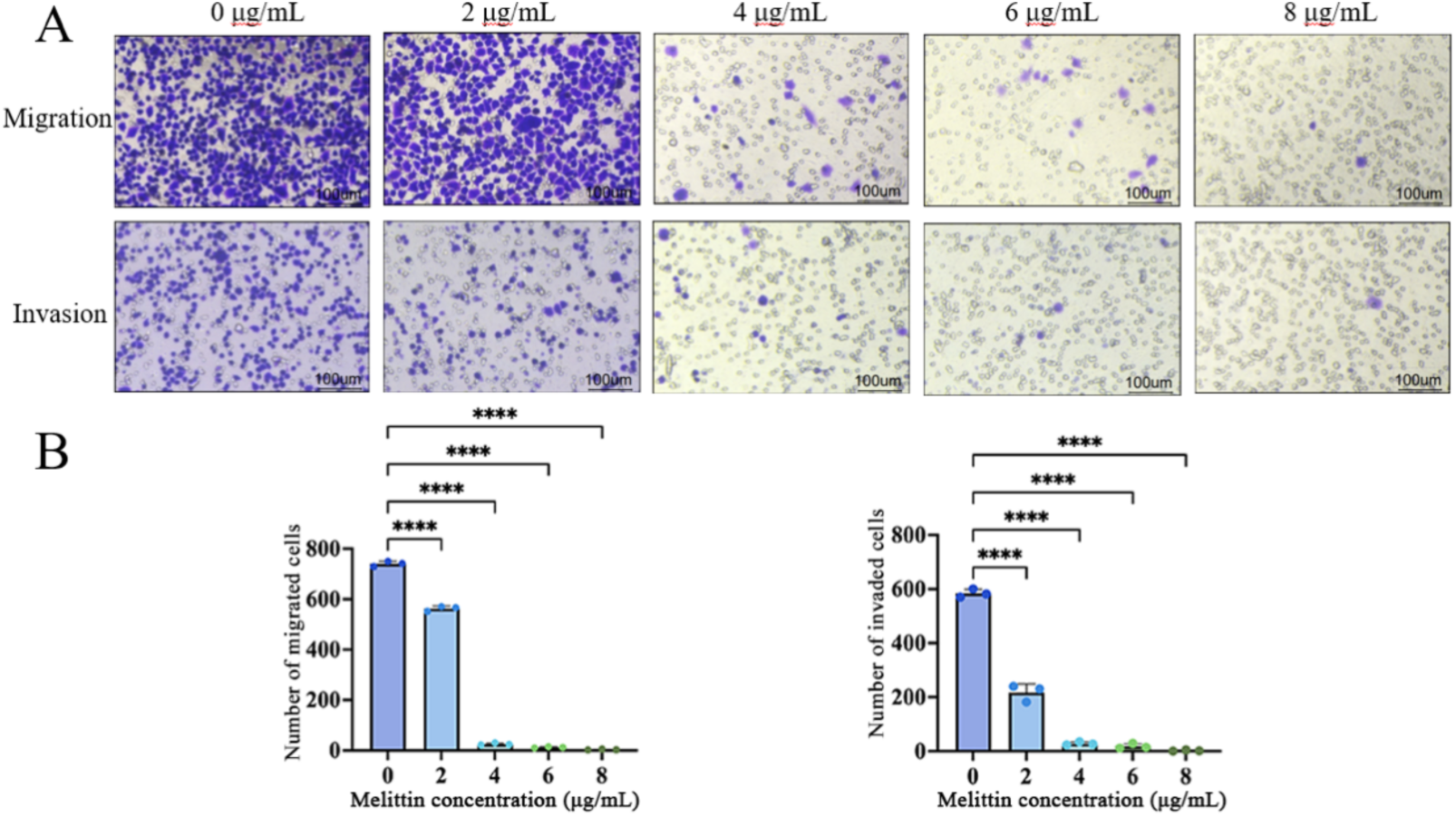
Melittin suppresses the invasion and migration of U14 cervical cancer cells in a dose-dependent manner. (A. Representative images of Transwell invasion assays of U14 cervical cancer cells treated with melittin (0, 2, 4, 6, and 8 µg/mL) for 24 h. B. Representative images of Transwell migration assays of U14 cervical cancer cells treated with melittin (0, 2, 4, 6, and 8 µg/mL) for 24 h. Quantitative data are presented as Mean ± SD from three independent experiments (n = 3). Statistical analysis was performed using one-way analysis of variance (one-way ANOVA) followed by Dunnett’s multiple comparisons test (vs. the 0 µg/mL C group). \**P* < 0.05, \*\**P* < 0.01, \*\*\**P* < 0.001, \*\*\*\**P* < 0.0001; ns, not significant.)

### 3.2 Melittin leads to death of U14 cervical cancer cells

To evaluate the cytotoxic effect of melittin on cervical cancer cells, U14 cervical cancer cells were treated with increasing concentrations of melittin (0, 2, 4, 6, and 8 µg/mL) for 24 h and subjected to PI/Hoechst double staining. Hoechst 33342 stained the nuclei of all cells (blue), whereas propidium iodide (PI) selectively labeled cells with compromised plasma membrane integrity (red) (Figure 2A). Compared with the untreated C group, melittin treatment markedly increased the number of PI-positive cells, indicating enhanced cell death (Figure 2A). Quantitative analysis (Figure 2B) suggested that the PI/Hoechst ratio increased with melittin concentration from 2 to 6 µg/mL, with the greatest increase observed at 4 µg/mL (\*\**P* < 0.0001 vs. control). In contrast, the PI/Hoechst ratio was reduced at 8 µg/mL and was not significantly different from that of the control group (ns). These findings demonstrate that melittin-induced cytotoxicity was concentration-dependent within the lower concentration range but did not further increase at the highest concentration tested.

**Figure 2.**
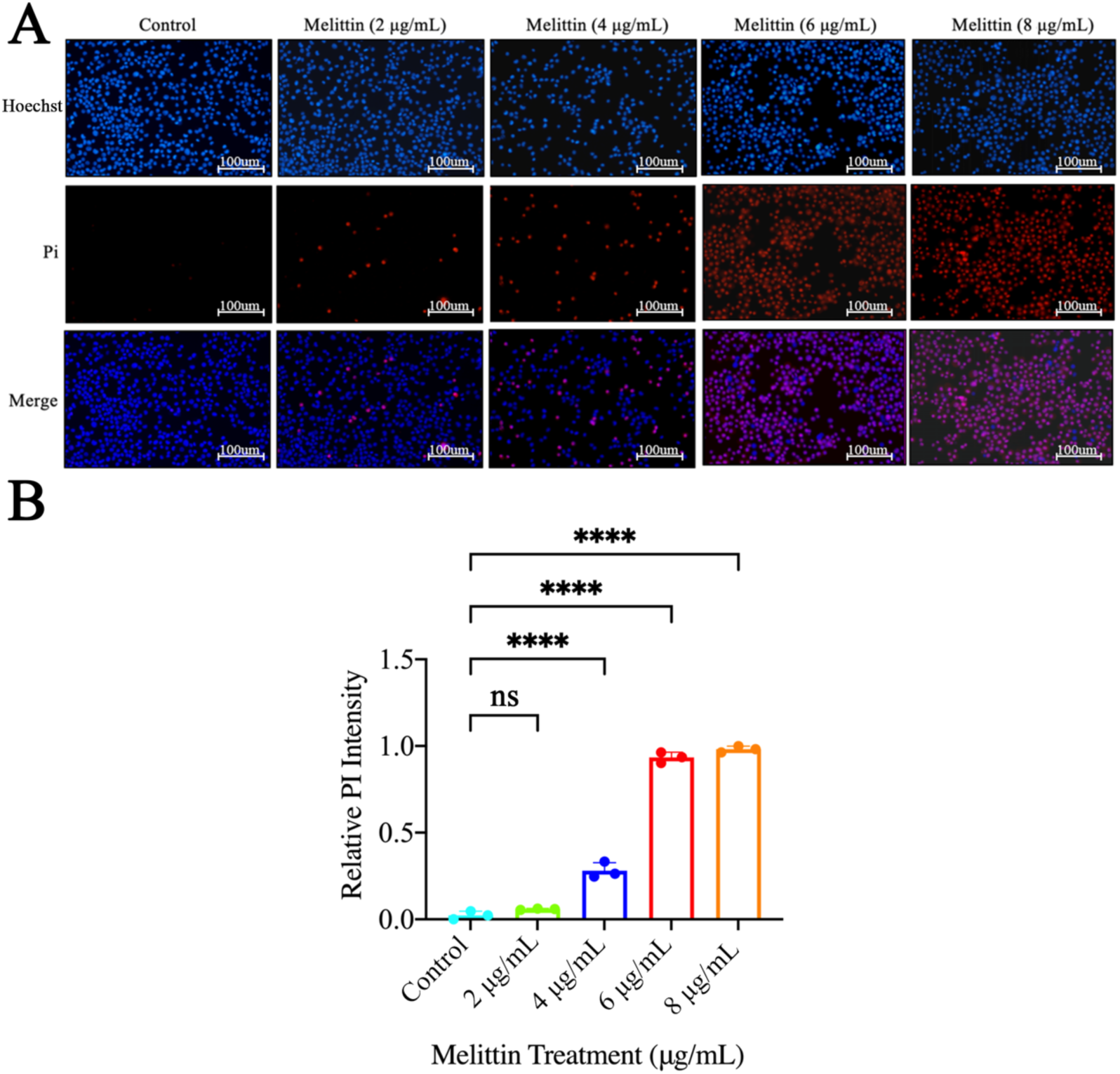
Melittin induces membrane damage and cell death in U14 cervical cancer cells. (A) Representative images of PI/Hoechst staining after 20 min treatment with melittin (0–8 µg/mL). Hoechst (blue) labels all nuclei; PI (red) labels damaged cells. Scale bar = 100 µm. (B) Quantification of the PI/Hoechst ratio. Data are shown as mean ± SD (n = 3). \*\*\*\**P* < 0.0001 vs. control; ns, not significant.

### 3.3 Transcriptional validation of oxidative stress-associated genes following mitochondrial oxidative phosphorylation remodeling

To further examine the oxidative stress-related changes identified by proteomic analysis, the transcriptional expression levels of representative redox-associated genes were determined by qRT-PCR. As shown in Figure 3, treatment altered the mRNA expression of genes involved in ROS production, antioxidant defense, and glutathione (GSH) metabolism.

**Figure 3.**
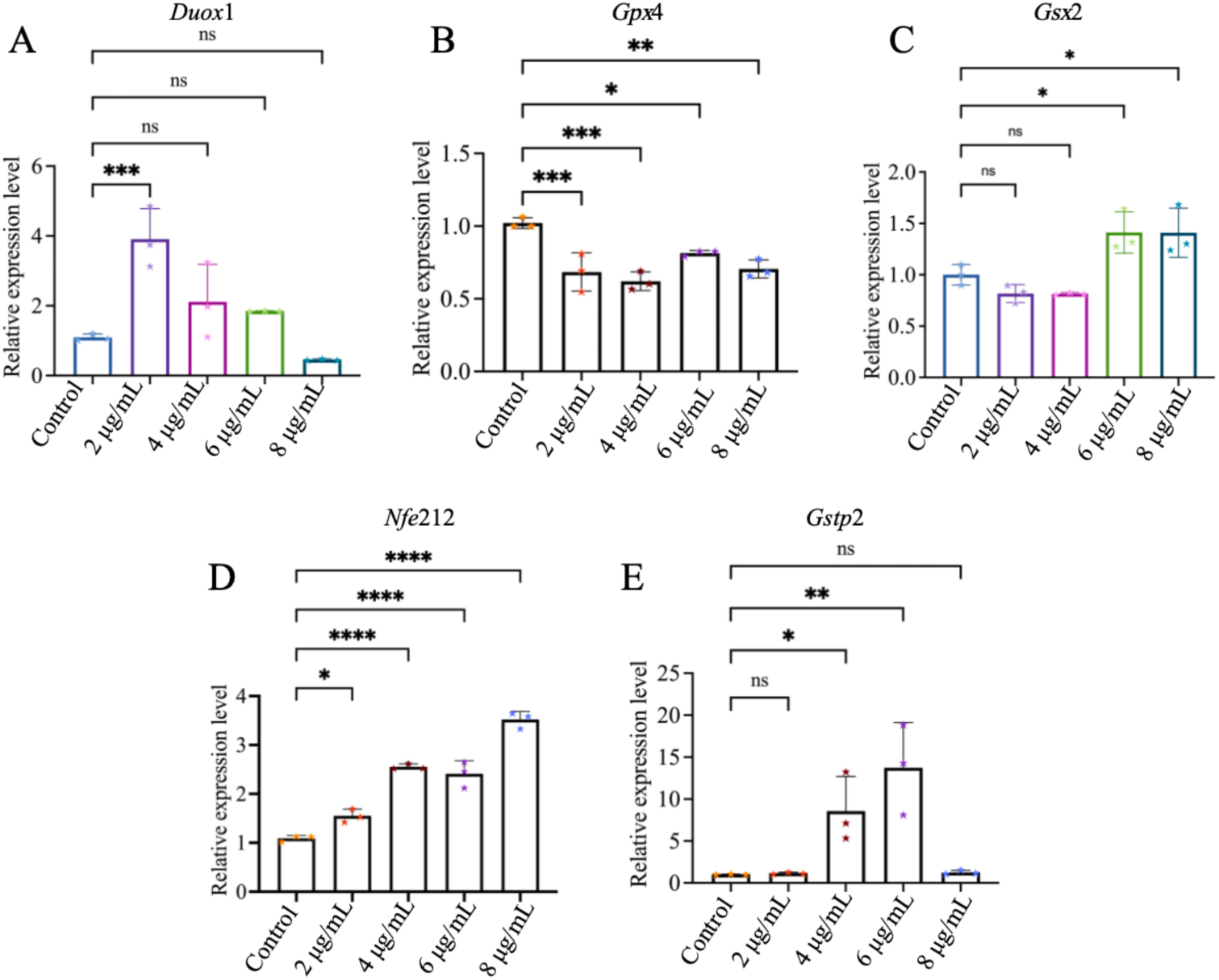
qRT-PCR validation of oxidative stress-related genes during mitochondrial OXPHOS remodeling. Expression levels of *Duox*1 (A), *Gpx*4 (B), *Gsx*2 (C), *Nfe*2l2 (D), and *Gstp*2 (E) were analyzed by qRT-PCR. Data are shown as mean ± SD (n = 3).

The expression of *Duox*1 (Figure 3A) changed following treatment and exhibited a dose-dependent trend. The mRNA level of *Gpx*4 (Figure 3B) was significantly decreased compared with the C group. In addition, the expression of *Gsx*2 (Figure 3C) was significantly altered after treatment. The transcript level of *Nfe*2l2 (Figure 3D) increased in a dose-dependent manner. Similarly, *Gstp*2 (Figure 3E) expression was significantly upregulated in the treatment groups compared with the C group.

Overall, qRT-PCR analysis demonstrated significant changes in the transcriptional expression of several oxidative stress-related genes following treatment, including *Duox*1, *Gpx*4, *Gsx*2, *Nfe*2l2, and *Gstp*2.

### 3.4 Quality control of DIA quantitative proteomics data

Pearson correlation analysis demonstrated high reproducibility among biological replicates in both the C and T groups, with correlation coefficients exceeding 0.98 (Figure 4A), indicating excellent sample consistency. DIA-based quantitative proteomic analysis was performed with a false discovery rate (FDR) of ≤1%, resulting in the identification of 163,087 peptides corresponding to 9,782 protein groups. The number of quantified proteins in individual samples ranged from 9,157 to 9,261, demonstrating a high depth of proteome coverage (Figure 4B, see also Table 1). Based on the screening criteria of P < 0.05 and |log₂FC| > 0.263, a total of 187 differentially expressed proteins (DEPs) were identified, including 71 upregulated and 116 downregulated proteins. GO enrichment score analysis further revealed a negative enrichment pattern of OXPHOS -related proteins in the T group (ES = −0.435, NES = −1.469, P = 0.014; Figure 4C), suggesting that this biological process was significantly affected following melittin treatment.

**Figure 4.**
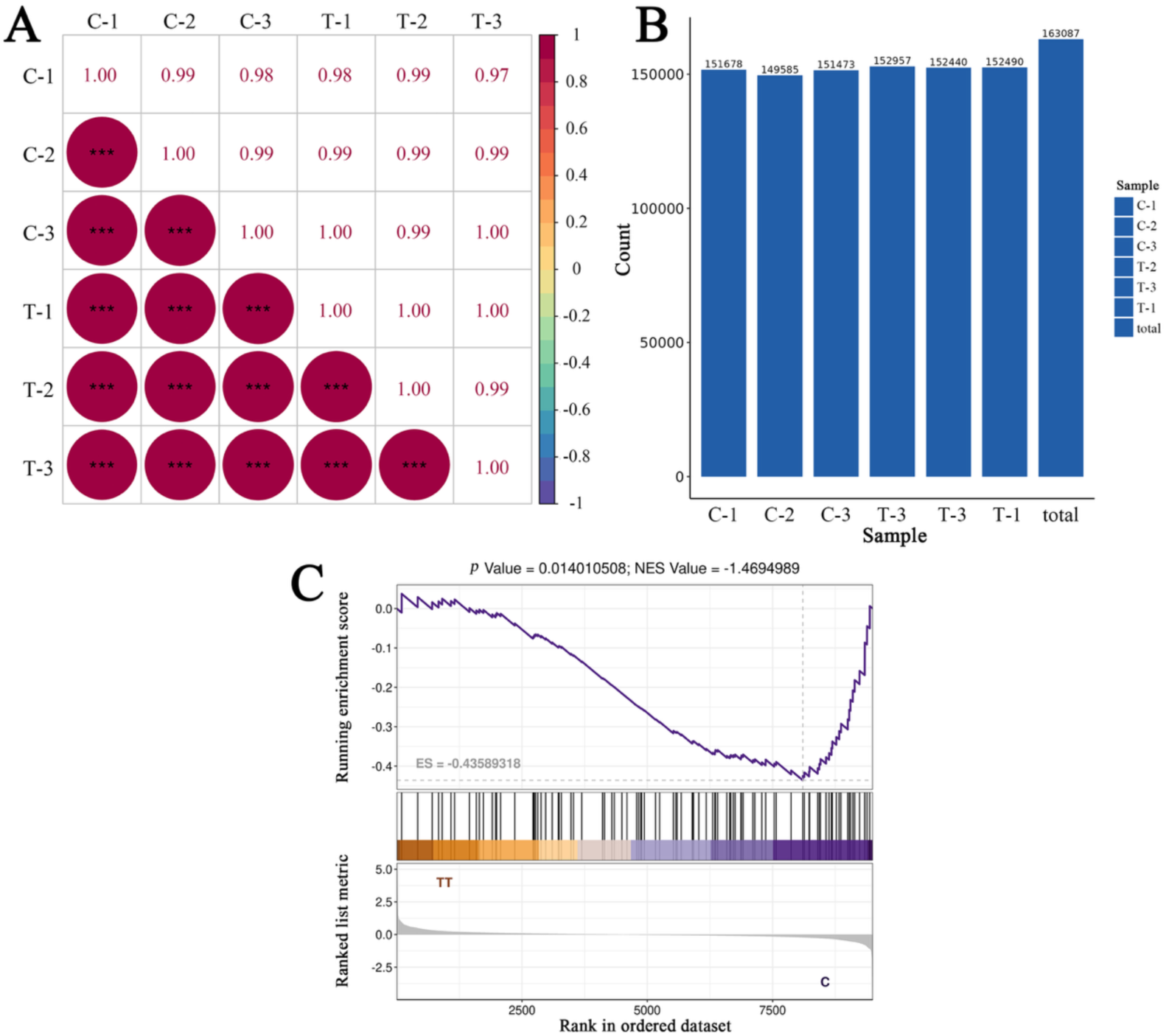
Quality control and proteomic profiling of U14 cervical cancer cells following melittin treatment. (A) Pearson correlation heatmap of biological replicates in control (C1–C3) and melittin treated groups (T1–T3). All intra group coefficients exceeded 0.98. (B) Total identified peptides (163,087) and quantified proteins per sample (9,157–9,261). (C) GSEA enrichment plot showing downregulation of the OXPHOS pathway (ES = –0.435, NES = –1.469, P = 0.014).

**Table 1.**
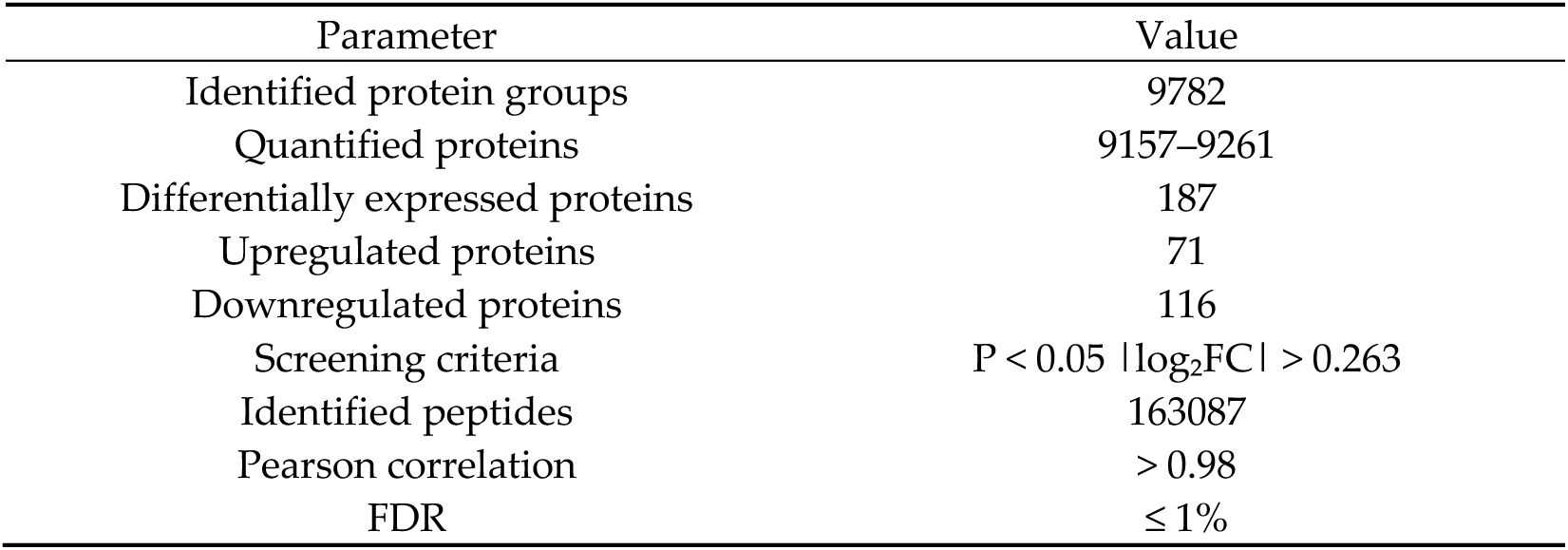
Summary of Astral-DIA proteomics data.

### 3.5 Identification and investigation of DEPs

DIA based quantitative proteomics identified a total of 9,782 protein groups, with individual samples yielding 9,157 to 9,261 quantified proteins, indicative of high reproducibility and consistency across biological replicates (Figure 5A). Differential expression analysis screened 187 DEPs in U14 cervical cancer cells following melittin treatment, comprising 71 up- and 116 down-regulated proteins (Figure 5B, see also Table S1). To gain further insight into the proteomic alterations induced by melittin, hierarchical clustering was performed on selected DEPs, the heatmap exhibited clear segregation between C and T groups (Figure 5C). Given that GSEA identified OXPHOS as a significantly downregulated pathway (Figure 4C), we further examined the expression patterns of OXPHOS related proteins. As shown in Figure 5C, the majority of OXPHOS associated proteins were downregulated in the T group, corroborating the GSEA findings and reinforcing the notion that melittin suppresses mitochondrial energy metabolism. Heatmap analysis of the top 100 DEPs revealed a clear separation between C and T groups, with the majority of proteins downregulated upon treatment, including multiple OXPHOS-related components (Ndufa12, Uqcrh, Cox6a1, Cox6b1), while Pnma8a and Myo1d were among the few upregulated proteins (Figure 5D).

**Figure 5.**
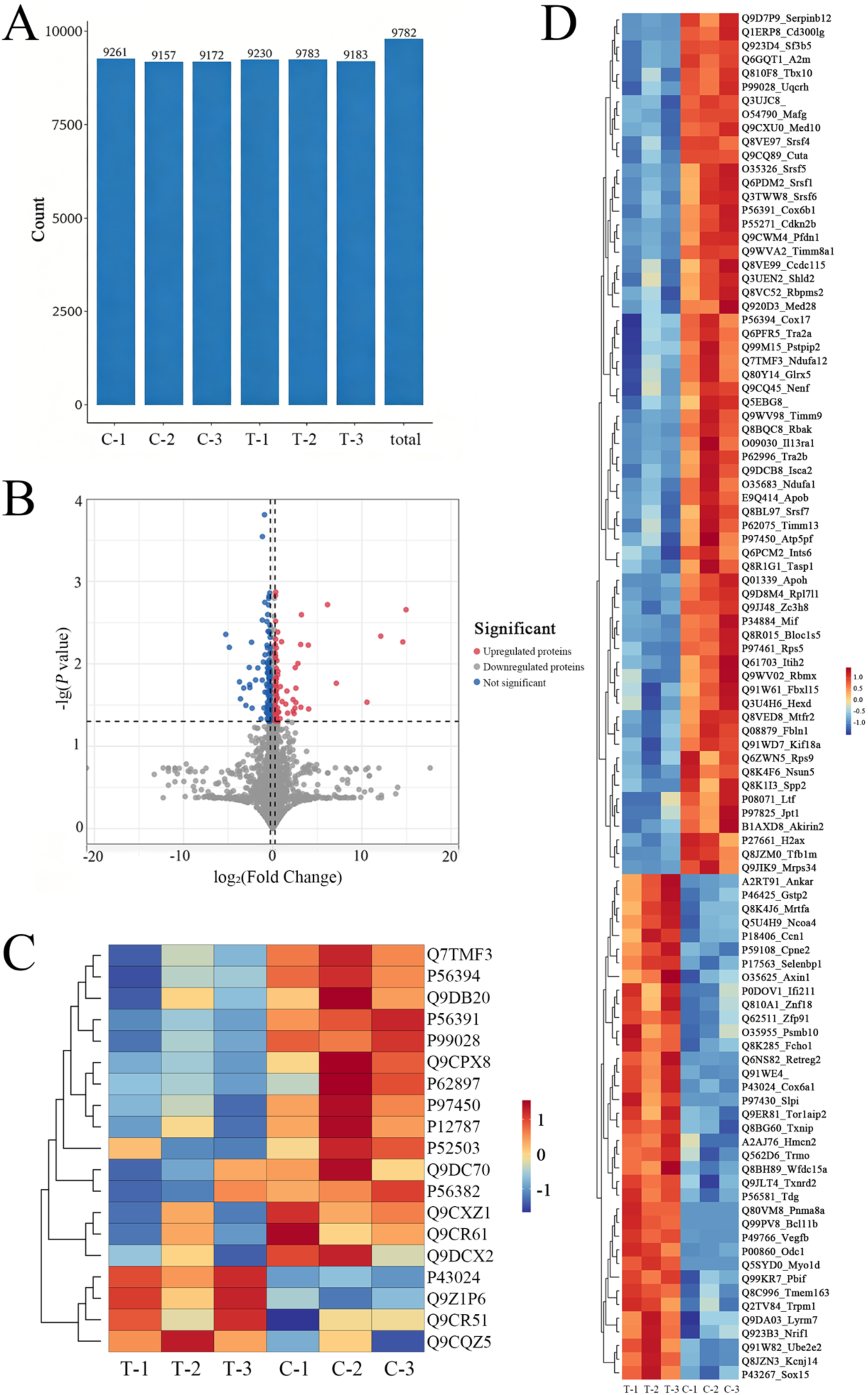
Proteomic profiling and differentially expressed proteins in melittin treated U14 cervical cancer cells. (A) Number of quantified proteins per sample and total identified protein groups. (B) Volcano plot showing 187 DEPs (71 upregulated, 116 downregulated) based on |log₂FC| > 0.263 and *P* < 0.05. (C) Heatmap of OXPHOS related proteins. (D) Heatmap of differentially expressed proteins between C and T groups.

### 3.6 GO term and KEGG pathway enrichment analysis of DEPs

KEGG pathway enrichment analysis suggested that OXPHOS exhibited the most significant enrichment, highlighting mitochondrial energy metabolism as a major target of melittin-mediated regulation (Figure 6A). In addition, several redox-associated pathways, including GSH metabolism, ferroptosis-related pathways, and chemical carcinogenesis–reactive oxygen species (ROS) signaling, were significantly enriched, suggesting that melittin treatment may perturb cellular redox homeostasis. Moreover, enrichment of the mitophagy pathway indicated potential alterations in mitochondrial quality control processes in response to mitochondrial stress.

**Figure 6.**
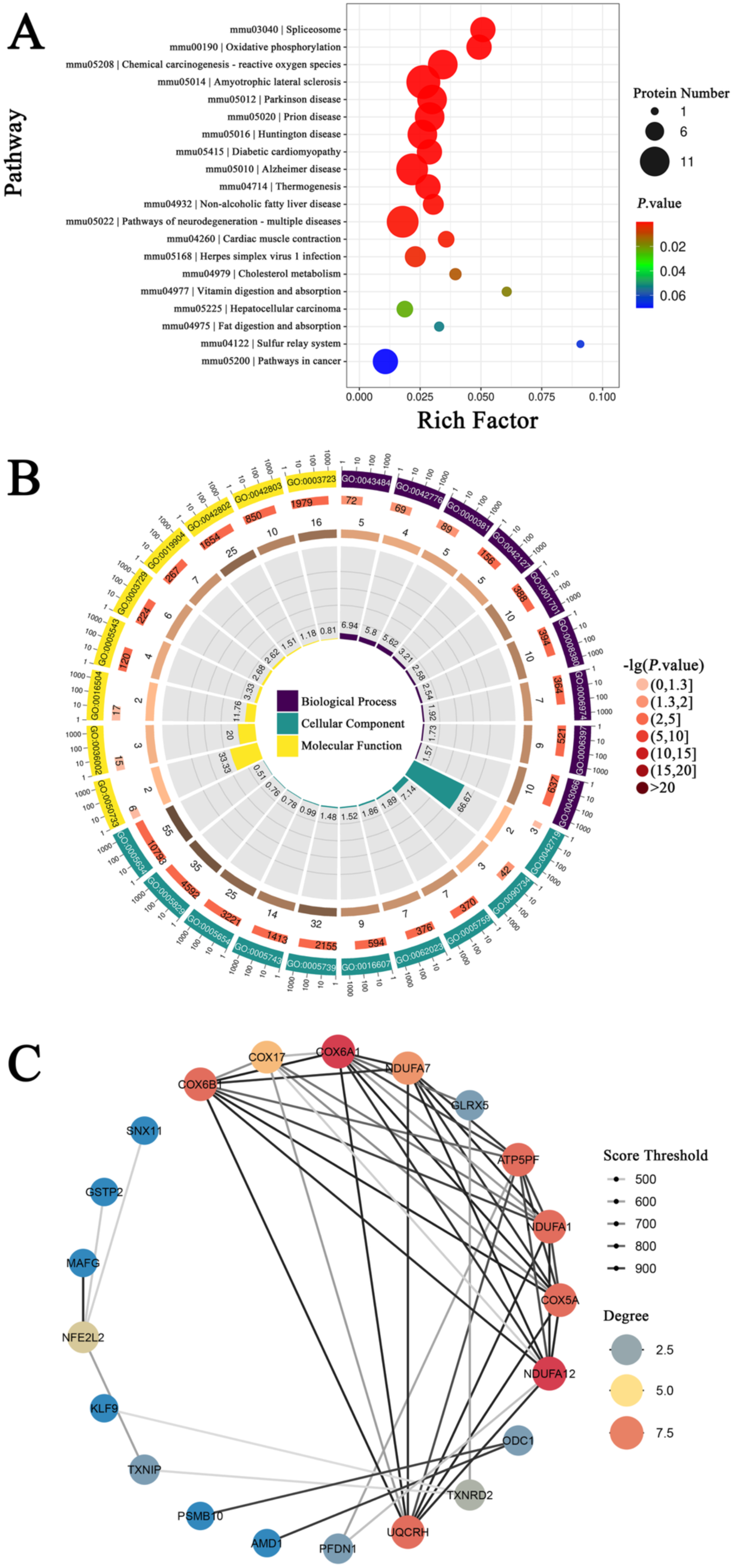
Functional enrichment and protein-protein interaction network analysis of differentially expressed proteins in melittin treated U14 cervical cancer cells. (A) Bubble diagram of KEGG pathways enriched by DEPs. (B) Loop diagram of GO terms enriched by DEPs. (C) PPI network of DEPs.

GO term enrichment analysis demonstrated that the identified DEPs were mainly associated with mitochondrial-related cellular components and biological processes involved in electron transport, oxidoreductase activity, and energy metabolism (Figure 6B). To further investigate functional interactions among the altered proteins, a PPI network was constructed, in which glutathione S-transferase pi 2 (GSTP2) was identified as representative hub proteins associated with redox regulation and metabolic adaptation. The differential expression of GSTP2 suggests that melittin induces coordinated alterations in antioxidant capacity and metabolic homeostasis (Figure 6C).

### 3.7 Melittin impairs mitochondrial oxidative phosphorylation and ATP production

Consistent with the KEGG pathway enrichment results, mapping of DEPs onto the OXPHOS pathway demonstrated that multiple proteins associated with mitochondrial respiratory chain complexes I–V were altered following melittin treatment (Figure 7A), indicating extensive remodeling of the mitochondrial oxidative phosphorylation machinery. To functionally validate these proteomic alterations, intracellular ATP levels were measured. As shown in Figure 7B, melittin treatment significantly reduced ATP production in a dose-dependent manner, with ATP levels progressively declining as the melittin concentration increased.

**Figure 7.**
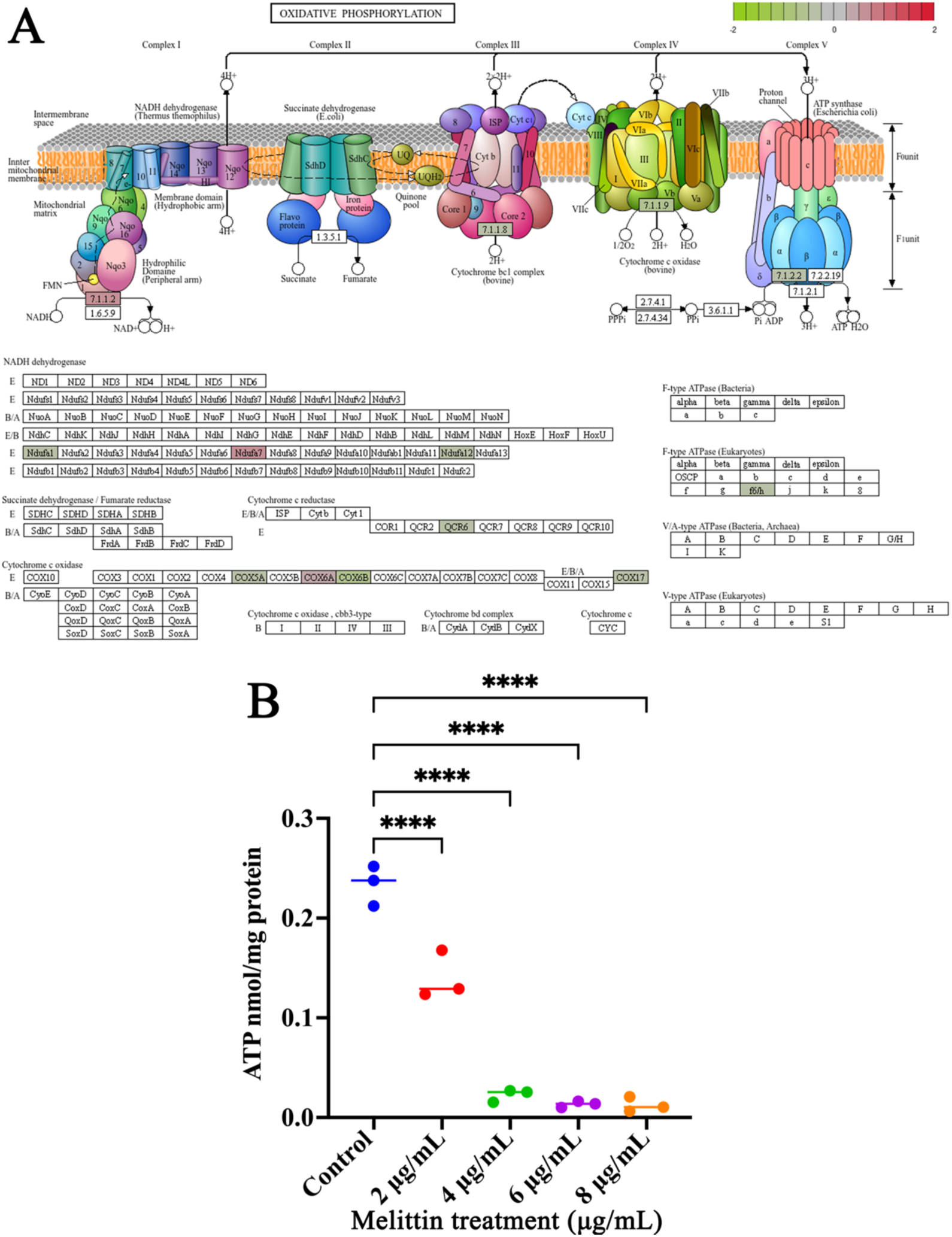
Melittin impairs mitochondrial oxidative phosphorylation and reduces ATP production in U14 cervical cancer cells. (A) Overview of the oxidative phosphorylation pathway. (B) ATP levels in U14 cervical cancer cells treated with different concentrations of melittin. Data are presented as mean ± SD. \*\*\*\**P* < 0.0001.

### 3.8 Western blot detection of GSTP2 in melittin-treated U14 cervical cancer cells

Western blot analysis was performed to validate the differential expression of GSTP2 identified by proteomic analysis. Compared with the control group, melittin treatment resulted in an increased GSTP2 protein level, as confirmed by densitometric quantification normalized to β-Tubulin (Figure 8). These results further support the involvement of GSTP2-mediated redox regulation in the cellular response to melittin.

**Figure 8.**
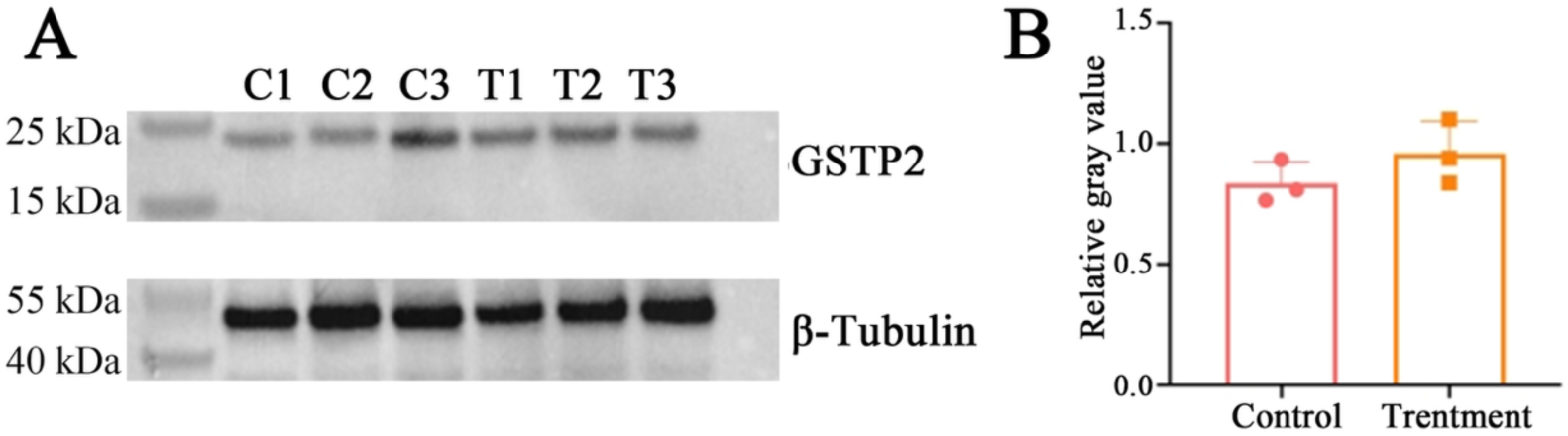
Validation of GSTP2 protein expression by western blot analysis following melittin treatment. (A) Western blot detection of GSTP2 protein expression in U14 cervical cancer cells treated with melittin. β-Tubulin was used as the loading control. (B) Densitometric quantification of GSTP2 expression normalized to β-Tubulin. C1–C3, Un-treated control groups, T1–T3, melittin-treated groups.

## 4. Discussion

Previous studies have shown that melittin suppresses the viability, migration, and invasion of multiple cancer cell types [33,34]. Consistent with these findings, our previous work showed that melittin significantly inhibited the proliferation [30,31], migration, and invasion of U14 cervical cancer cells in a dose-dependent manner. In the present study, Transwell assays further confirmed the inhibitory effects of melittin on cell migration and invasion (Figure 1), indicating that melittin effectively suppresses the malignant phenotype of cervical cancer cells. In addition, PI/Hoechst staining demonstrated a concentra-tion-dependent increase in PI-positive cells following melittin treatment (Figure 2), suggestive of enhanced cell death. These findings are in agreement with previous studies reporting the cytotoxic activity of melittin against various cancer cell types.

To further validate the oxidative stress-associated alterations revealed by proteomic profiling, the transcriptional expression patterns of key redox-related genes were examined using qRT-PCR. Melittin exposure induced significant changes in the expression of genes involved in ROS regulation and GSH metabolism (Figure 3). Specifically, *Duox*1 (Figure 3A) and *Gsx*2 (Figure 3C) exhibited distinct expression alterations, indicating perturbation of redox homeostasis following melittin treatment. Moreover, the marked downregulation of *Gpx*4 (Figure 3B) suggested compromised antioxidant defense capacity and enhanced vulnerability to oxidative stress. Conversely, *Nfe*2l2 (Figure 3D) and its downstream target *Gstp*2 (Figure 3E) were significantly upregulated, reflecting the activation of the NRF2-dependent antioxidant response as an adaptive cellular mechanism. Nevertheless, this compensatory response appeared insufficient to counteract the excessive oxidative burden induced by melittin, thereby contributing to redox imbalance and subsequent cell death. Collectively, these findings corroborate the proteomic observations and suggest that mitochondrial energy metabolism and GSH-dependent redox regulation may represent critical molecular events underlying the anticancer effects of melittin.

Here, DIA-based quantitative proteomic profiling provided comprehensive coverage of the cellular proteome and revealed extensive molecular alterations in response to melittin treatment (Figure 4B). Functional enrichment analysis demonstrated a significant negative enrichment of oxidative phosphorylation-associated proteins following melittin exposure (Figure 4C), highlighting mitochondrial energy metabolism as a major target of melittin-induced cellular perturbation. As the central organelle responsible for maintaining cellular bioenergetics, metabolic homeostasis, and redox regulation, mitochondria play a critical role in cancer cell survival and adaptation. Dysregulation of mitochondrial oxidative phosphorylation has been increasingly recognized as a hallmark of cancer metabolism and a potential vulnerability for therapeutic intervention [35]. Therefore, the observed suppression of oxidative phosphorylation-related proteins suggests that melittin may disrupt mitochondrial metabolic reprogramming, thereby impairing the energetic demands required for tumor cell maintenance and survival. These findings provide novel mechanistic insights into the antitumor activity of melittin in cervical cancer and highlight mitochondrial metabolic dysfunction as a potential contributor to its anticancer effects.

DIA-based quantitative proteomic analysis revealed extensive proteomic remodeling in response to melittin treatment, with 187 differentially expressed proteins identified in U14 cervical cancer cells (Figure 5A–B). Hierarchical clustering analysis demonstrated a distinct separation between control and melittin-treated groups (Figure 5C), indicating substantial alterations in the cellular proteomic landscape following melittin exposure. Consistent with the GSEA results (Figure 4C), a predominant downregulation pattern was observed among OXPHOS-associated proteins, including critical mitochondrial respiratory chain components such as Ndufa12, Uqcrh, Cox6a1, and Cox6b1 (Figure 5C–D). Mitochondrial dysfunction and metabolic reprogramming are recognized as fundamental characteristics of cancer biology, enabling tumor cells to adapt to increased energetic and biosynthetic demands [36]. However, the molecular mechanisms driving mitochondrial metabolic alterations during tumor progression remain incompletely understood [37]. Our findings suggest that melittin-induced suppression of multiple respiratory chain components may compromise mitochondrial electron transport efficiency and energy production, thereby perturbing mitochondrial homeostasis in cervical cancer cells. Collectively, these results identify mitochondrial oxidative phosphorylation remodeling as a potential molecular basis underlying the antitumor activity of melittin.

KEGG pathway enrichment analysis identified OXPHOS as the most significantly altered pathway following melittin treatment (Figure 6A), indicating that mitochondrial energy metabolism represents a major molecular process affected by melittin exposure. Increasing evidence suggests that mitochondrial metabolic reprogramming and redox imbalance are closely associated with cancer progression and therapeutic responses. Ferroptosis, a distinct form of regulated cell death characterized by iron-dependent lipid peroxidation, is tightly regulated by cellular antioxidant systems and metabolic pathways, particularly GSH-dependent redox regulation [38]. In the present study, the enrichment of GSH metabolism, ferroptosis-associated pathways, and ROS-related signaling pathways suggests that melittin treatment induces profound alterations in cellular redox regulation.

Consistently, GO enrichment analysis revealed that melittin-responsive proteins were predominantly associated with mitochondrial components, electron transport processes, oxidoreductase activity, and energy metabolism (Figure 6B), further supporting mitochondrial metabolic remodeling as a major cellular response to melittin. Moreover, PPI network analysis identified GSTP2 as a potential hub protein involved in redox regulation and metabolic adaptation (Figure 6C), suggesting the activation of compensatory antioxidant responses in response to melittin-induced oxidative stress.

Mitochondria function as essential regulators of cellular bioenergetics and are also major sources of intracellular ROS [39]. Impaired electron transport chain activity can promote ROS accumulation, lipid peroxidation, and oxidative damage, thereby contributing to redox imbalance and cell death signaling [40]. Meanwhile, GSH, the most abundant intracellular antioxidant molecule, plays a central role in maintaining redox homeostasis, and disruption of GSH metabolism has been implicated in oxidative stress and ferroptosis regulation [41]. Therefore, the coordinated enrichment of mitochondrial metabolic pathways and GSH-related processes observed in this study suggests that melittin may exert its antitumor activity by perturbing mitochondrial bioenergetic function and compromising cellular redox homeostasis. Collectively, these findings highlight the interplay between mitochondrial metabolic disruption and oxidative stress regulation as potential mechanisms underlying melittin-mediated anticancer effects.

To further validate the impact of melittin on mitochondrial oxidative phosphorylation, the altered proteins were mapped onto the OXPHOS pathway. Multiple components of mitochondrial respiratory chain complexes I–V were affected following melittin treatment (Figure 7A), confirming extensive remodeling of the oxidative phosphorylation machinery. Consistently, functional assessment revealed a significant reduction in intracellular ATP production in a dose-dependent manner (Figure 7B), demonstrating that melit-tin-induced OXPHOS disruption directly impairs mitochondrial energy metabolism.

Notably, among the identified DEPs, GSTP2 is closely associated with GSH metabolism and intracellular redox regulation. GSTP2, a member of the glutathione S-transferase family, plays a vital part in antioxidant defense and ROS detoxification [42,43], Here, western blot analysis confirmed the upregulation of GSTP2 identified by proteomic analysis (Figure 8), further supporting the involvement of GSH-dependent antioxidant regulation in the cellular response to melittin. Together, these findings demonstrate that melittin suppresses mitochondrial energy metabolism while simultaneously activating redox adaptation mechanisms, highlighting the coordinated regulation of mitochondrial dysfunction and redox homeostasis during its anticancer activity.

**Figure 9.**
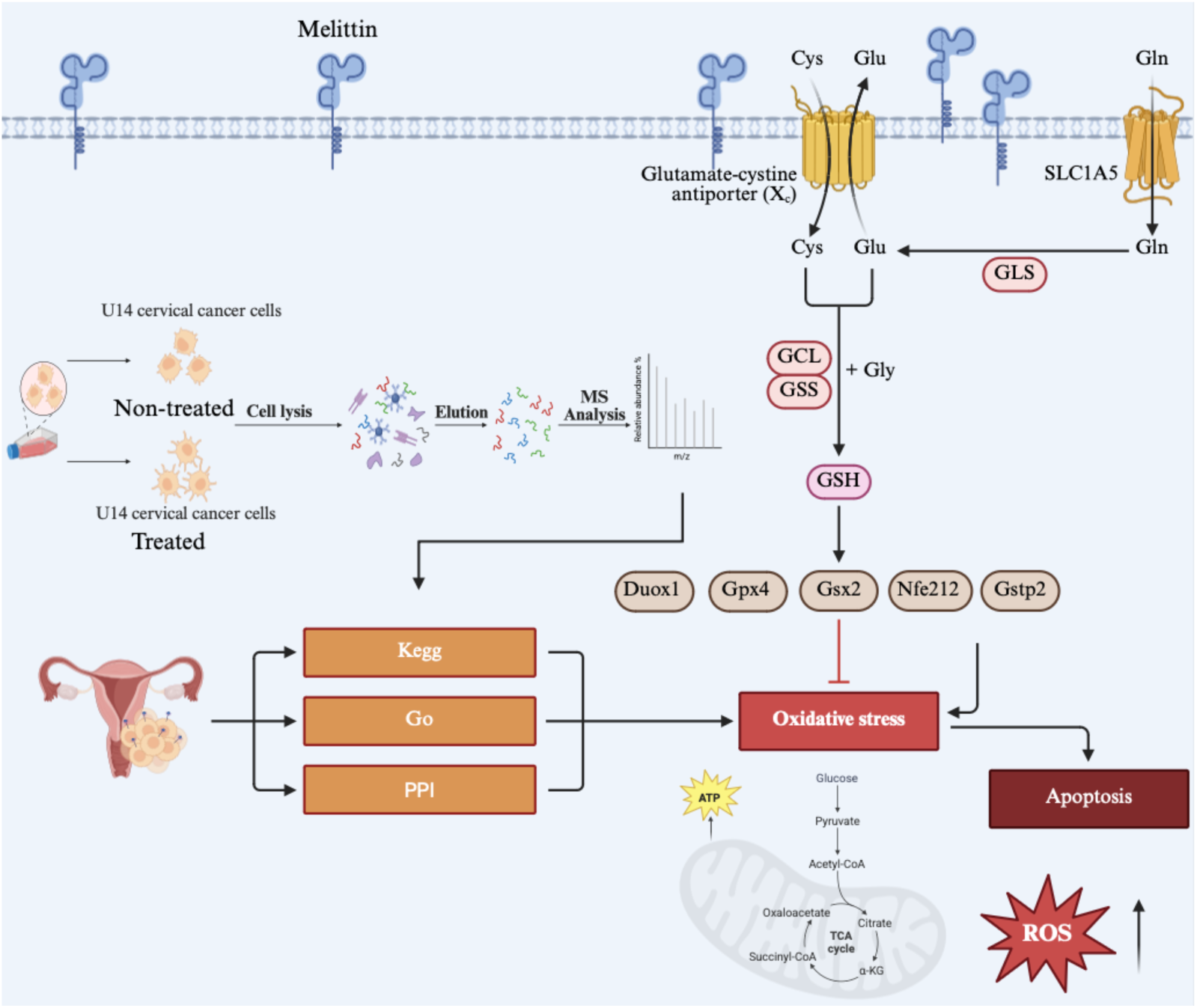
A schematic diagram of the effect of melittin on U14 cervical cancer cells.

## 5. Conclusions

In conclusion, melittin significantly suppresses the migration and invasion of U14 cervical cancer cells and regulates biological processes associated with mitochondrial function and redox homeostasis, with OXPHOS and GSH metabolism emerging as key pathways underlying its anticancer effects.

## Supplementary Materials

The following supporting information can be downloaded at Table S1

## Author Contributions

The following statements should be used “Conceptualization, R.Z. and M.W.; methodology, R.Z.; software, M.W.; validation, R.Z., M.W. and K.Z.; formal analysis, M.W.; investigation, K.Z.; resources, H.Z.; data curation, S.L.; writing—original draft preparation, M.W.; writing—review and editing, J.J.; visualization, T.Y.; supervision, J.Q., R.G. and D.C.; project administration, J.Q., R.G. and D.C.; funding acquisition, J.Q., R.G. and D.C. All authors have read and agreed to the published version of the manuscript.”

## Institutional Review Board Statement

Not applicable.

## Data Availability Statement

All the data are contained within the article.

## Acknowledgments

All of the authors thank all editors and reviewers for their constructive comments and recommendations. We express our sincere gratitude to the Biorender team for providing this invaluable resource to the scientific and educational communities. It has significantly enhanced the efficiency and quality of our work. All individuals included in this section have consented to the acknowledgement.

## Abbreviations

The following abbreviations are used in this manuscript:

DEPs: Differentially expressed proteins
GO: Gene Ontology
KEGG: Kyoto Encyclopedia of Genes and Genomes
PPI: Protein–protein interaction
Astral-DIA: Astral data-independent acquisition
OXPHOS: Oxidative phosphorylation
RT-qPCR: Reverse transcription quantitative PCR
PBS: Phosphate-buffered saline
MS: Mass spectrometry
GSEA: Gene set enrichment analysis
DO: Disease Ontology
FDR: False discovery rate
One-way ANOVA: One-way analysis of variance
PI: Propidium iodide
GSH: Glutathione
ROS: Reactive oxygen species

## Disclaimer/Publisher’s Note

The statements, opinions and data contained in all publications are solely those of the individual author(s) and contributor(s) and not of MDPI and/or the editor(s). MDPI and/or the editor(s) disclaim responsibility for any injury to people or property resulting from any ideas, methods, instructions or products referred to in the content.

